# The primacy model and the structure of olfactory space

**DOI:** 10.1101/255661

**Authors:** Hamza Giaffar, Sergey Shuvaev, Dmitry Rinberg, Alexei A. Koulakov

**Affiliations:** Cold Spring Harbor Laboratory, Cold Spring Harbor, NY 11724; Neuroscience Institute, New York University Langone Medical Center, New York, NY 10016; Department of Physics, New York University, New York, NY

## Abstract

Understanding sensory processing relies on establishing a consistent relationship between the stimulus space, its neural representation, and perceptual quality. In olfaction, the difficulty in establishing these links lies partly in the complexity of the underlying odor input space and perceptual responses. Based on the recently proposed primacy code for concentration invariant odor identity representation and a few assumptions, we have developed a theoretical framework for mapping the odor input space to the response properties of olfactory receptors. We analyze a geometrical structure containing odor representations in a multidimensional space of receptor affinities and describe its low-dimensional implementation, the primacy hull. We propose the implications of the primacy hull for the structure of feedforward connectivity in early olfactory networks. We test the predictions of our theory by comparing the existing receptor-ligand affinity and connectivity data obtained in the fruit fly olfactory system. We find that the Kenyon cells of the insect mushroom body integrate inputs from the high-affinity (primacy) sets of olfactory receptors in agreement with the primacy theory.

## INTRODUCTION

In color vision, our ability to predict the perceptual quality of color from a spectrum of incident light rests on a small number of receptor types at the neural periphery and also depends on our understanding of the properties of these receptors. The dimensionality of the color space is defined by the 3 types of receptors, i.e., 3 degrees of freedom, two of which define the planar coordinates of the color and one that computes the total intensity of the light. The olfactory system operates with a much larger number of receptor types at the sensory periphery (∼350 in humans, ∼1200 in rodents, and ∼60 in flies^1–5^); the mapping of the chemical stimulus space to the receptor and perceptual space remains an unresolved problem. The discovery of such a large family of olfactory receptors (ORs)^6^ has promoted the idea that the dimensionality, *D*, of this olfactory space is high and comparable to the number of OR types, *N*.

There is mounting evidence, however, that olfactory perceptual space is low dimensional^7, 8^. Low dimensional embeddings of human perceptual data and its fitting by a curved manifold of dimension *D* < 10, which accounts for >80% of variance in the data, suggests that the number of odorant parameters relevant to the human olfactory system is <10 (Refs.[^8–10^]). A recent success to predict olfactory metamers for humans using a mixture model, which describes each odorant with ∼20 parameters, also suggests low dimensionality of the odor perceptual space^11^. Insights into the structure and dimensionality of olfactory perceptual space should guide our understanding of information processing in this sensory system.

A number of features of early olfactory neural circuits are strongly conserved across diverse species, from insects to mammals^12, 13^. Olfactory sensory neurons (OSNs) expressing ORs of the same genetic type converge on their respective glomeruli in the olfactory bulb (OB) or antenna lobe (AL). To a first approximation, each glomerulus represents the average level of activation of a single OR type. This information is projected to several higher-level processing centers, such as the Piriform cortex (PCx) in mammals and the mushroom body (MB) in insect, by axons of second order projection neurons. The logic of connectivity between the OB/AL and these downstream target areas has been under intense scrutiny as it seems to convey the nature of features important for olfactory processing. The convergent evolution of this ‘canonical’ olfactory circuit may suggest a conserved logic of odor information processing across phyla^14^. If this is true, then it should be possible to construct a general theory of olfactory information processing relevant to a wide range of species.

As in the case of vision, where a color percept is formed from light with a broad spectrum of monochromatic waves, a majority of ethologically relevant odors are mixtures of many tens or hundreds of components^15, 16^. Such complex mixtures of volatile chemicals are usually thought to be perceived synthetically in humans, in the sense that a single ‘odor object’ is discerned for any complex mixture, as opposed to elementally, as a sum of parts^17^. In addition, the perceptual identity of an odor is generally stable over a range of stimulus and neural parameters, including concentration, background and noise in neural circuits^18–22^. This feature of the neural code may contribute to the ability of animals to identify sources of smells at varying distances.

How can an odorant maintain constant identity despite changes in concentration? As odorant concentration increases from low to high, in general, more OR types are becoming activated^23, 24^. The representations of odor identity in high and low concentration regimes can therefore be linked by a set of ORs activated at low concentration that is active in both regimes. This template comprised of high affinity OR types to a given odorant is called the *primacy set* and the model for concentration-invariant odor coding relying on primacy sets is called the *primacy model*^25^. Despite the apparent simplicity, the primacy model can explain many psychophysical phenomena of odor perception and is compatible with known olfactory neural network organization^25–28^.

Here, we propose a new theoretical framework for mapping olfactory chemical space to the neural spaces of OR/glomerular and cortical representation. Our theory is based on the following main assumptions: (i) The stimulus or odor space is of relatively low intrinsic dimensionality, *D*. (ii) The *primacy-coding hypothesis*: odor identity is encoded by a small number of OR types of highest affinity to a given odor. We will study the implications of these assumptions for the evolution of OR ensembles and develop a set of statistical methods to test these assumptions. We demonstrate these methods using recently published data on connectivity in the fly olfactory system.

## RESULTS

### Primacy coding model

We will consider a model in which the activation of a receptor *f_ro_*, as a function of odorant concentration, *c_o_*, depends only on one parameter *K_ro_*, the affinity of receptor *r* to odorant *o*, and can be described by the mass action law:

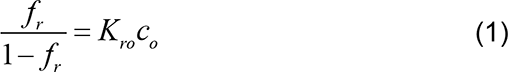

In logarithmic concentration coordinates, *f_r_* = *f_r_* (*c_o_*) is a logistic function (Fig. 1A)^24^. When the activation of the receptor reaches certain threshold, θ, the changes in the response can be detected by the downstream system, at which point, the OR becomes activated and can participate in the odorant coding. Our theoretical conclusions are not strongly affected by the choice of activation threshold of activation, as long as it is similar across the receptors. For the sake of simplicity, we will define an OR as active if its response to an odorant is higher than a half of the maximum activity level, i.e., *f_r_* > θ = 1/ 2, which corresponds to the odorant concentration *c_o_* > 1/ *K_ro_*. Importantly, in this model, receptors activated at the lowest concentration remain active at higher concentrations.

**Figure 1.**
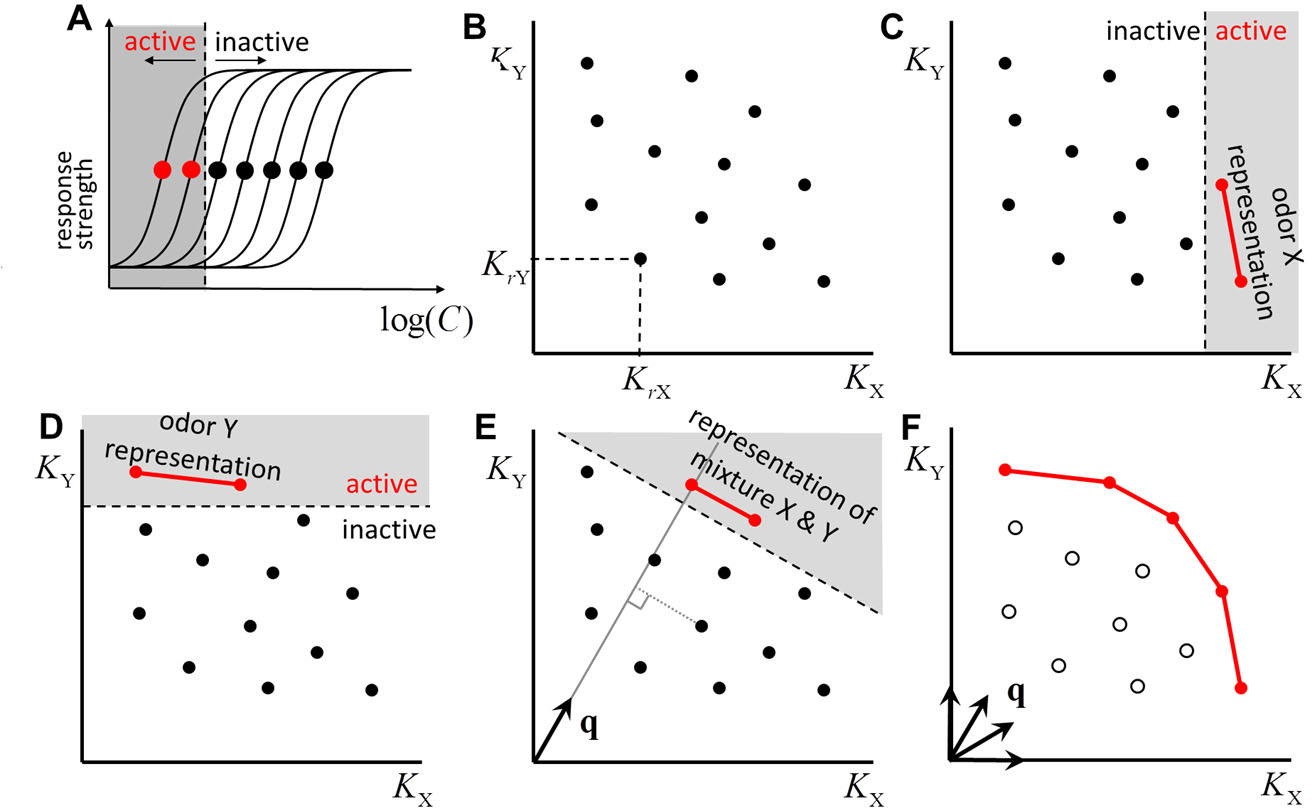
Primacy model in 2D. **A.** Receptor response curves as a function of odorant concentration. Solid circles correspond to effective threshold concentrations or inverse affinities: *c^th^* = (*K*)^−1^. For some concentration (dashed line), the ORs with a threshold below this concentration are in an active state: (red solid circles). **B.** Representation of receptors in 2D space of affinities for odorants *X* and *Y* : *K_X_* and *K_Y_*. **C.** An odorant *X* at concentration *c_X_* activates all receptors for which *K_rX_ c_X_* > 1 (red). For the given primacy number *p* = 2, the identity of odor *X* is defined by the two most sensitive receptors (red segment). **D.** The same for odor *Y*. **E.** A mixture of two odorants *X* and *Y* activates receptors, which are above a line perpendicular to a unit vector **q** = [*c_X_*, *c_Y_*] / *c*, and defined by an effective mixture concentrastion 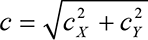. Primacy sets for all possible mixture, vectors **q**, define a *primacy hull* (red), 1D line in 2D space. Receptors in the primacy hull (red) are retained in the genome. All other ORs (empty circles) and eventually eliminated from the genome (pseudogenized).

According to the primacy coding hypothesis, the identities of a few most sensitive OR types determine the perceptual odor identity associated with a stimulus. We define the primacy set for a given odorant as the set of *p* OR types activated by the odorant at the lowest concentration (Fig. 1A). Different odorants evoke activity in different sets of most sensitive receptors. In principle, the primacy number, *p*, could vary across odorants; however, for simplicity, here we will assume it to be fixed.

### A two-dimensional odor space

To explore the implications of primacy coding for the organization of OR ensemble, we consider an odor space comprised of two odorants (*X*, *Y*) and their mixtures. In this case, each OR type can be represented as a point in a 2D space of affinities for the two odorants with coordinates **K***_r_* = (*K_rX_*, *K_kY_*). An example arrangement of a receptor ensemble in a 2D odor space is shown in Fig. 1B. Introducing a pure odorant *X* at a given concentration, *c_X_* partitions the space into two half spaces; one in which receptors are active, *K_rX_ c_X_* 1, and the other in which they are inactive, *K_rX_ c_X_* < 1 (Fig. 1C). Increasing *c_X_* moves the boundary between active and inactive receptors and expands the zone of active receptors from high to low *K_X_*. If we consider a primacy model in which two glomeruli are required to identify an odor (a *p* = 2 primacy model), there is a concentration of odor *X* at which the first two glomeruli are activated. These glomeruli represent OR types of highest affinity to odorant *X* and represent the identity of *X* in primacy coding mechanism.

Similarly, the primacy set corresponding to an odor *Y* can be assigned by introducing pure odor *X* at a low concentration and expanding the active zone by increasing the concentration until the first *p* = 2 OR types are activated (Fig. 1D).

To consider mixtures, we extend the equation **Error! Reference source not found.**to the case of two odorants. Assuming the independence of individual odorant-receptor interactions, we have (Supplementary Methods S):

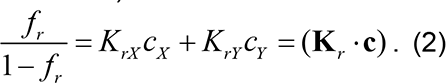

Here (**K***_r_* ·**c**) is the scalar product between the vector of affinities of a receptor *r* for the set of molecules *X* and *Y*, **K** *_r_* = [*K_rX_*, *K_rY_*], and the vector of concentrations **c** = [*C_X_*, *C_Y_*]. Receptor activations can be described by a similar equation that explicitly accounts for the overall concentration of the odor mixture:

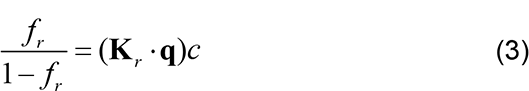

Here (K_r_*c)* is the Euclidian length of the concentration vector representing the overall mixture concentration and **q** = [*c_X_*, *c_Y_*] / *c* is a unit vector in the direction defined by the ratio between the mixture components. For a given mixture concentration *c*, the activation level of an OR is determined by the projection of its affinity vector **K***_r_* and the unit vector of mixture components **q**, i.e. **K***_r_* ·**q**. The most active ORs are those with the largest value of **K***_r_* ·**q** (Fig. 1E). For a *p* = 2 primacy model, the primary receptors are the two points with the largest projection onto the vector **q**, i.e. the two ORs activated at the lowest mixture concentration (Fig. 1E). By considering the primacy sets of all possible mixtures, i.e. all possible **q** s in this 2D odor space, we trace out a hull that contains all OR types that belong to at least one primary set (Fig. 1F). We call this structure a ***primacy hull***.

The primacy hull contains all OR types that belong to at least one primacy set. According to the primacy model defined here, odor identity is encoded by the OR types of highest affinity to a given odorant, i.e., primary ORs. ORs that have low affinity to any odor and are not members of any primacy set and do not participate in odor identity coding. Unless such ORs participate in encoding odorant features other than odor identity or are used for other non-olfactory coding roles^14^, they are likely to be pseudogenized and eliminated from the genome (Fig. 1F). Thus, one of the predictions of the primacy model is that the functional ORs should belong to a primacy set of at least one odorant and therefore belong to the primacy hull, unless they have some other function.

Eqs. (1-3) follow from the receptor-ligand binding model which does not include a number of important mechanisms involved in odorant-OR interactions, such as antagonism^29, 30^ or potential multiple odorant binding sites per OR^31^. Nevertheless, following^12, 31–36^, we will adopt this model for mathematical convenience. We consider the implications of nonlinear interactions between odorant molecules for the primacy model in Supplemental Information S5.

### Higher-dimensional odor spaces

As in the preceding 2D model, in case of more odorants present, ORs can be represented as vectors of affinities to individual odorants in a D-dimensional odor space: **K** *_r_* = [*K_r_*_1_, *K_r_* _2_,…, *K_rD_*]. The receptor response to a mixture is described by Equation (2) with the odor mixture represented by the unit vector: **q** =[*c*, *c*,…, *c*] / *c* where *c* = (*c*^2^_1_ + *c*^2^_2_ + … + *c*^2^*_D_*)^1/2^. At a given concentration, a *D* − 1-dimensional plane orthogonal to the vector **q** separates the active and inactive receptors. Increasing an odor concentration moves this plane toward the origin and recruits additional receptors. The first *p* activated ORs form a primacy set for the mixture defined by the vector **q**. The primacy number, *p*, and the dimensionality of the odor space, *D*, are independent parameters. In the previous example we explored the case in which *p* = 2; each odor identity is represented by two nodes (OR types), which may be thought of as a segment. In general, a primacy set can be represented by a geometric object called a (*p* −1) -simplex: *p* = 2 corresponds to a line segment, *p* = 3 to a triangle, and *p* = 4 to a tetrahedron (Fig. 2A).

**Figure 2.**
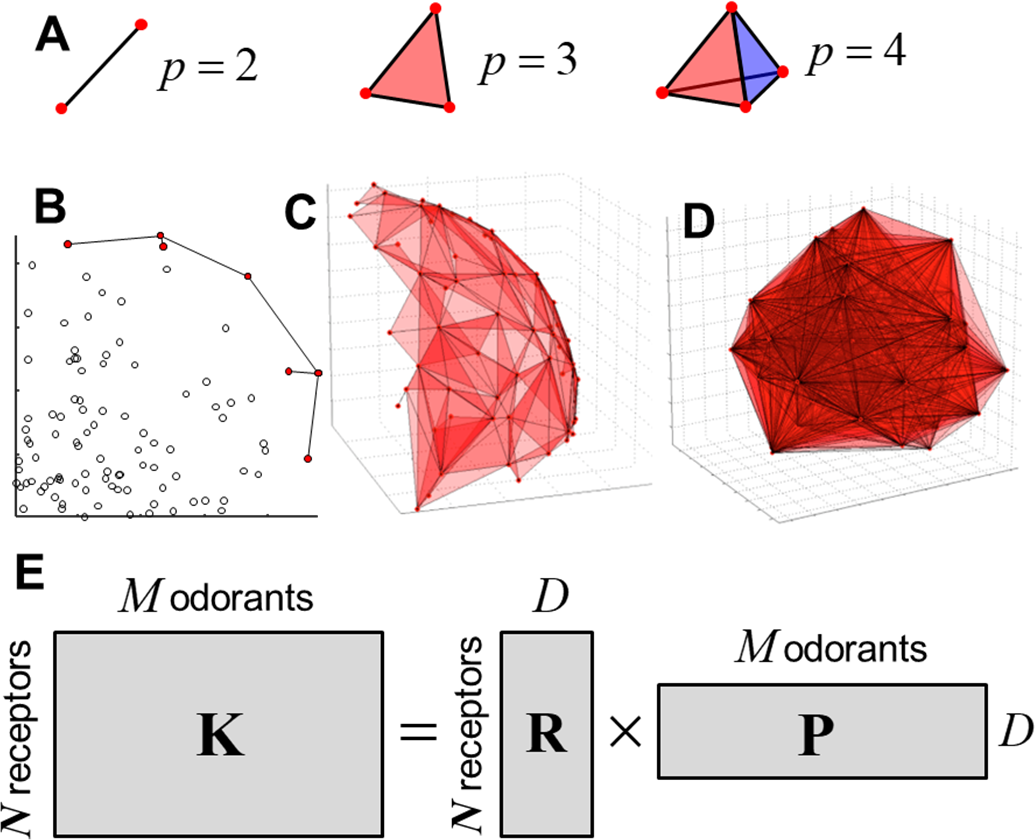
Primacy hull in higher dimensions. **A.** Geometrical representation of primacy sets for different primacy numbers: *p* = 2: 1-simplex, a segment; *p* = 3: 2-simplex, a triangle; *p* = 4 : 3-simplex, tetrahedron. **B**. An example of primacy hull for a random set of points in 2D (*p* = 2). A primacy hull is a set of simplexes that reside on the extremes of the given set of points. **C.** A primacy hull for *D* = 3, *p* = 3. A ∼2D surface is tessellated by triangles, each of them representing an independent odor identity. **D.** The same for *D* = 6, *p* = 7. 6D manifold is projected onto 3D space for visualization. **E.** Decomposition of affinity matrix ***K*** into two low-dimensional matrices: ***P*** is a low-dimensional matrix of basic odor features, and ***R*** is a matrix of effective receptor affinities for these basic features.

The primacy hull is a simplicial complex (a collection of simplexes) obtained by sweeping planes through a collection of points; for each plane, the first *p* encountered points (those of highest affinity) are associated into a simplex and added to the complex (Fig. 1). The primacy hull therefore includes the convex hull as a subset, as well as additional points internal to the convex hull (Fig. 2B). Examples of primacy hulls for *D* = 3, *p* = 3 and for *D* = 6, *p* = 7 are shown in Fig. 2B-D. The latter hull is projected onto 3D for visualization.

The full set of volatile molecules likely includes millions of compounds, yet the description of a primacy hull in *D* ∼ 10^6^ is not very useful. As discussed in the introduction, some experimental evidence suggests that the dimensionality of the odor space is low. We will next formulate the low-dimensionality assumption within our framework. Equation (2)**Error! Reference source not found.** relates receptor activity to the receptor’s response properties, i.e. affinities to different odorants and concentrations of mixture components. Affinities can be determined as inverse concentration thresholds 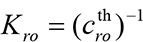 estimated from concentration dependencies of neural responses (Fig. 1A). Here 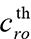 is a concentration of a monomolecular odorant *o*, at which a receptor *r* is activated at half of its maximum magnitude.

Affinities *^K^ro* used in Equation (2) describe the affinity of *N* ORs to *M* odorants. These quantities can be combined into an large affinity matrix **K** which has dimensions *N* × *M* (*N* ∼ 10^3^, *M* ∼10^7^). If **K** can be accurately represented by a product of two matrices of much smaller size, **R***_N_*_×_*_D_* and **^P^***D*×*M*, where *D* = *N*, *M*, we can say that the dimensionality of the odor space is small (Fig. 2E):

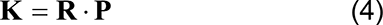

Here **P** is a *D* × *M* matrix placing every molecule into a D-dimensional space of “properties of interest” to the olfactory system. It describes to what degree these properties are present in each of the *M* molecules. These properties may include molecular weight, measures of polarity, size, and other potentially more complex molecular features^8,10, 37^ or their combinations. **R** is a *N* × *D* matrix representing every receptor as a point in the D-dimensional space of molecular properties. The affinity of the receptor *r* for a (monomolecular) odorant *o* is determined by the scalar product of the corresponding row in the matrix **R** and a column in the matrix **P**, both of which are D-dimensional. If the number of odorant parameters sampled by the olfactory system is on the order of the dimensionality of human perceptual space, i.e. ∼ *D* ∼ 10, then the simplification resulting from Equation (4) may be substantial^8,10^.

Equation (4) describes the affinities of receptors for pure odorants. Responses to mixtures can be obtained by combining equations (2) and (4), which results in

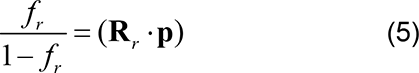

Here, we introduced a D-dimensional vector **p** = **Pc** describing the concentration of relevant properties in the odorant mixture described by the vector **c**. For example, if the ORs were sensitive to the molecular weights of monomolecular compounds, as suggested in Ref. [^10^], one component of the vector **p** would represent the concentration of molecular weight, i.e. the molecular weight per liter of gas. The role of the *D* × *M* property matrix **P** is therefore to project the concentration vectors of mixture components which may have millions of components to a much smaller *D* -dimensional space of properties relevant to the olfactory system. The vector **R***_r_* is a *D* -dimensional row of the matrix **R** which describes the OR sensitivity to these properties. The activation level of receptor by the mixture is determined by the dot product between the small-dimensional vectors of affinities and property concentrations, similarly to Equation (2). Thus, the approach described above, including the geometric constructs, such as primacy sets and the primacy hull, is valid in the case of low-dimensional olfactory space [Equation (4)] despite odorant mixtures including, potentially, millions of monomolecular components. In this case, instead of the description in the space of concentration vectors, the primacy sets and the primacy hull are built in the space of relevant odorant properties. Overall, we suggest that in case of low-dimensional odor space, the mechanisms of coding of odor identity in the primacy sets described above applies in the space of odorant properties.

### Connectivity to higher brain regions

The coactivation of primary ORs could be detected by feedforward connectivity between ORs/glomeruli and higher processing centers, if the connectivity contained information about the primacy sets. In this mechanism, the projections from individual primacy sets of ORs/glomeruli would converge on distinct cells in the piriform cortex (PC) or the mushroom body (MB) in insects (Fig. 3). PC neurons are expected to respond when the corresponding primacy ORs are co-activated (Fig. 3A, B). This prediction suggests that, although feedforward connectivity may appear to lack any obvious structure, it may contain high-order correlations induced by the presence of the primacy hull in responses. It also suggests that the connectivity is related to the OR responses – PC/MB cells integrate inputs from OB/AL neurons with the strongest affinity for an odorant. We will later test this prediction using recent Al-to-MB connectivity and OR-odorant affinity data from *D. Melanogaster*.

**Figure 3.**
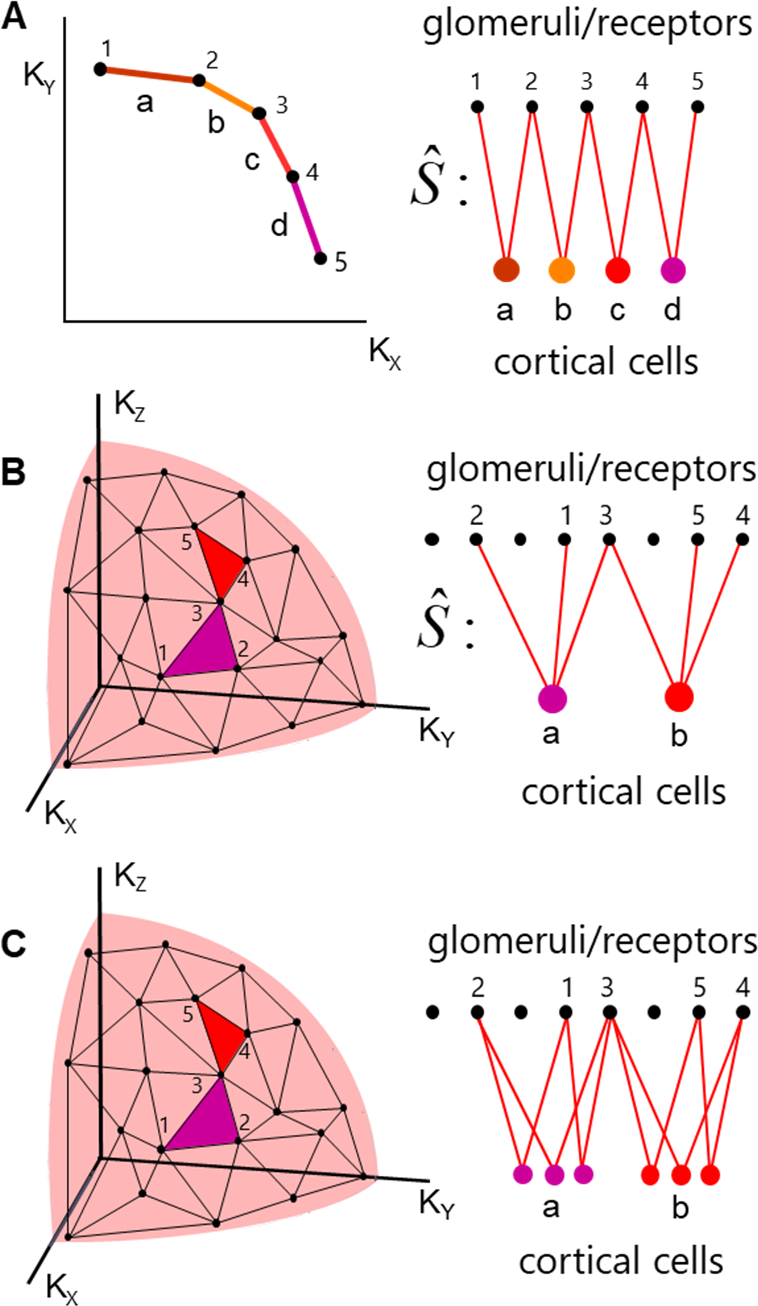
Suggested feedforward circuit which can process the primacy information. **A**. *Left:* Primacy hull for *D* = 2 and *p* = 2, a, b, c, d are discriminable odor identities. *Right*: connectivity between ORs 1, 2, …5 and cortical cells (insect mushroom body cells) corresponding to different odor identities. The glomeruli from the same primacy sets converge to the same cortical cells. **B.** The same for *D* = 3 and *p* = 3. Only two example simplexes are illustrated. **C.** Subprime connectivity model. Individual cortical cells represent the faces of primacy simplexes (sides of the triangles). Individual odor identities are encoded by populations of neurons marked ‘a’ and ‘b’.

In the connectivity model proposed above (Fig. 3A, B), only a single PC neuron responds to a presentation of an odorant, corresponding to a primacy simplex. In contrast, experimental data in mice shows that the PC contains about *N* ≍ 2 ·10^5^ pyramidal neurons out of which about 5%, i.e. about *f* ∼ 10^4^ respond to an odorant^38, 39^. One way in which a primacy model can generate 10^4^ responses is to assume that individual neurons in the PC represent faces of the primacy simplex rather than the full (*p*— 1)-simplex itself (Fig. 3C). These faces are themselves simplexes. For example, a triangle or a 2- simplex contains three sides as faces, which are also 1-simplexes. A tetrahedron, a 3-simplex, contains four triangles (2-simplexes) as faces (Fig. 2A), etc. In general, the number of *n*-point faces of a primacy (*p* −1) -simplex is given by the binomial equation

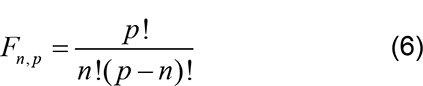

For example, a 4-vertex simplex (tetrahedron) contains 4!/3!/(4-3)!=4 3-vertex faces (triangles) and 4!/1!/(4-1)!=6 2-vertex faces (segments, Fig. 2A).

We propose that responses of cells in higher brain regions such as PC or MB reflect the activations of simplexes that are subsets or faces of the primary simplex corresponding to the presented odorant’s identity. This may explain the large population of cells activated in PC and MB in response to an odor. For example, in mouse PC, an odor activates *f* ∼ 10^4^ cells ^38, 39^, which roughly corresponds to the primacy number is *p* =16, and the number of converging connections onto on PC cell *n* = 8 : *F* ≍ 1.3·10^4^. We will further call the faces of the primary simplex the *subprime* simplexes. The complete set of subprime simplexes of a given degree uniquely represents the primary simplex. For example, in Fig. 2A, the set of four triangles uniquely represents the primary tetrahedron. Such a coding schema may provide some robustness to noise. A distributed representation of an odor is much less sensitive to failures of individual neurons to be activated. If a single neuron in the PC or MB represents an odorant identity (Fig. 3A, B), silencing this neuron will eliminate the perception of this smell. In the case when individual neurons represent subprime simplexes corresponding to the same odor (Fig. 3C), failure of activation of individual neurons due to the presence of noise can be compensated by a pattern completion mechanism implemented by associative circuits in PC^40^. This can be accomplished if subprime simplexes corresponding to the same primary simplex are connected by synapses with positive strength, similarly to a Hopfield network. Overall, we suggest that cells activated in the PC represent the faces of the primary simplex corresponding to the stimulus identity. These representations can be created by feedforward OB-to-PC (or AL-to-MB in insects) connectivity that contains primacy hull structure in the weight matrix and may be facilitated by the recurrent connectivity in the PC.

### Experimental predictions

According to our model, the AL-MB or OB-PC connectivity is expected to contain a distributed representation of the primacy hull. Specifically, we would expect connectivity data to i) have a low-dimensional component that is consistent across members of a species and ii) individual MB/PC neurons integrate inputs from high affinity ORs to an odorant (primacy sets).

Below we test these two predictions in the fruit fly (*D. Melanogaster*) using 2 independent connectivity datasets from two individual animals and the OR-odorant affinity data. We will use OR and glomerulus interchangeably to refer to a genetically defined olfactory information channel consisting of a single glomerulus and its homotypic ORs.

### Using existing data on connectivity and OR affinities to test the primacy model

In the fly, the OR activity is relayed to the MB via projection neurons (PNs); a majority of PNs receive input from only a single glomerular channel. Each OR/glomerular channel is associated with several PNs. The principal cells of the MB, the Kenyon cells (KCs), are the major target of PN axons, with a single KC integrating ∼5-6 Glomeruli/OR type inputs. Two recent studies describe PN-KC connectivity in two adult flies. The FlyEM dataset contains PNs originating from all 51 olfactory glomeruli and terminating at a large number of KCs (N_KC_=1784 out of a total estimated ∼2000-2500 KCs per MB hemisphere)^41^. The FAFB dataset includes connectivity for 51 glomeruli and 1344 KCs^42^. Using this binarized connectivity data (Fig. 4A and B), we can interrogate the logic governing KC integration of OR channels.

**Figure 4.**
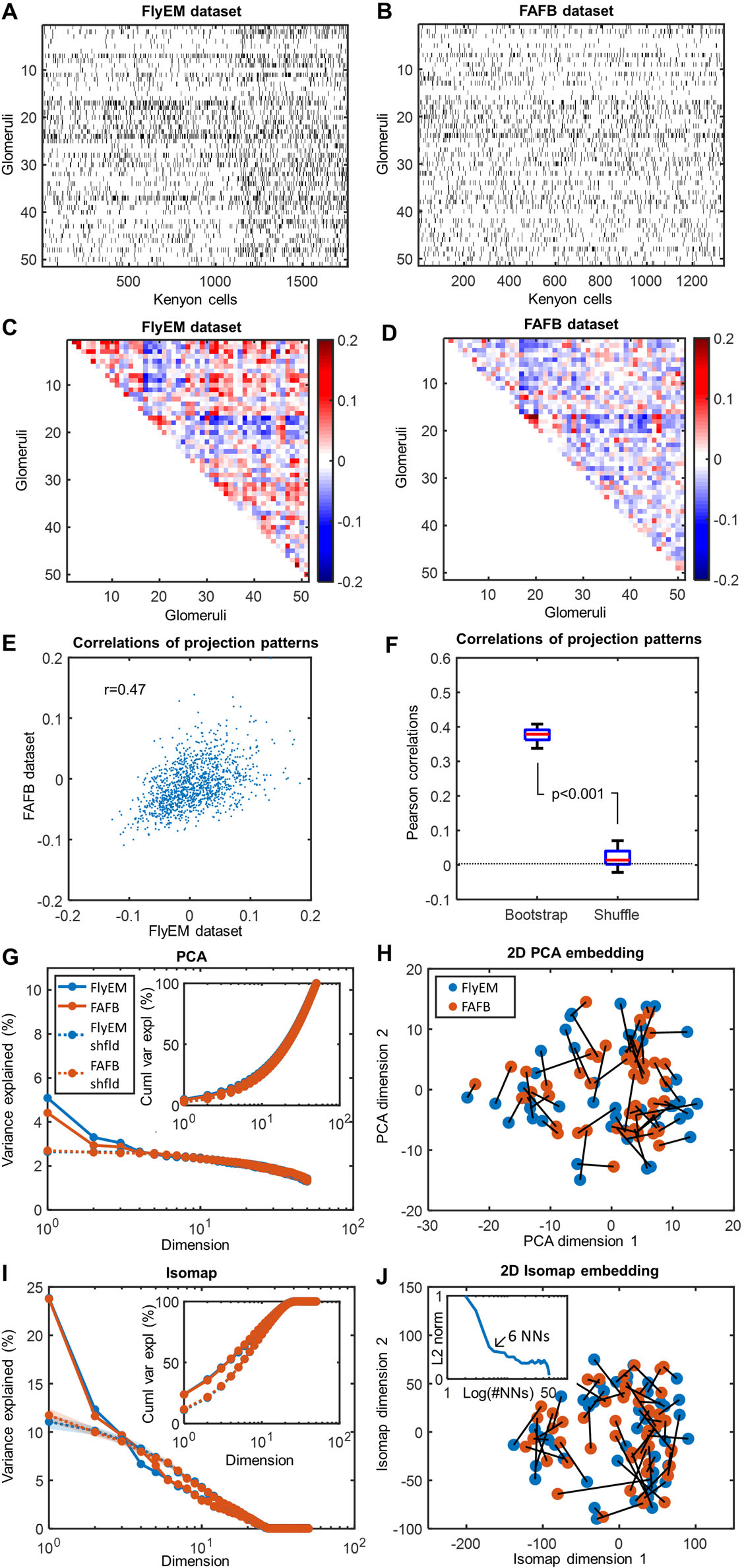
Non-random low-dimensional structure in uPN-KC connectivity that is conserved across animals. **A**, **B**. Glomerulus- KC connectivity matrices from FlyEM and FAFB datasets. **C, D**: Glomerulus-Glomerulus connectivity similarities (Pearson correlations of connectivities). **E**. Glomerulus-Glomerulus connectivity similarities in two datasets against each other. The correlation in Glomerulus-Glomerulus connectivity similarities is r=0.5 (p<0.01). **F**. Similarity between datasets disappears if one of the datasets is shuffled while preserving the connectivity matrix in- and out- degrees (right)^43^. **G**. Variance explained per dimension as a function of the PCA dimension (Inset – total cumulative variance explained). PCA analysis shows that two linear dimensions are significantly different from random. **H**. Connectivity matrices in two datasets projected onto the first two dimensions. Points represent individual glomeruli. The same glomeruli in two datasets (different animals) are connected by black segments. The same glomeruli in different datasets reside near each other in the 2D embedding suggesting that the first two dimensions of the connectivity matrix are conserved across datasets. **I, J**. The same analysis using a non-linear low-dimensional embedding technique (Isomap) shows that the first two dimensions in the connectivity data are both different from random (**I**), explain more variance in the data than the linear algorithm (PCA), and are conserved across datasets (**J**).

To test the prediction that the PN-KC data contains a low-dimensional component that is consistent across individual flies (Prediction i), we first compare the FlyEM and FAFB connectivity matrices. We aligned the two data matrices along the PN dimension by grouping PNs based on the identity of the glomeruli/ORs that they receive the primary input from. As there is no simple way to align KC identities across different animals, we started with analyzing the *similarity* of connectivity between individual glomeruli. These similarities were defined as Pearson correlation coefficients between pairs of glomeruli in terms of their projections to KCs computed for both FlyEM and FAFB datasets (Fig. 4C, D). We observed a substantial correlation between the similarity matrices (R=0.50, Fig. 4E). The correlation was eliminated by shuffling either connectivity matrix (Fig. 4F). Following Ref. [^43^], we used a shuffling procedure that preserves the number of connections from each glomerulus to KCs and from each KC to glomeruli (KC in- and glomerulus out-degrees). The presence of a significant correlation in connectivity between the two animals suggests that glomerulus-KC connectivity matrices share a common structure between the two individual animals.

Above we hypothesized that ORs sample a low-dimensional subspace of the affinity space, this subspace should be reflected in the connectivity structure consistent across members of the same species (Hypothesis i). To further characterize this common structure in the two connectivity datasets, we applied linear and nonlinear dimensionality reduction techniques to glomeruli-glomerulus similarity matrices. Linear dimensionality reduction methods, such as the principal component analysis (PCA), place the objects (here – glomeruli/ORs) on a flat surface, while nonlinear methods, such as Isomap, arrange them on a curved manifold of a given dimension. The quality of these embeddings can be assessed by the amount of variance in data explained by the embeddings. We applied both methods to glomerulus-glomerulus connectivity similarity matrices (Fig. 4C, D). First, we applied the PCA (flat) method (Fig. 4G, H). We found that the first two dimensions of the PCA space explain more variance than the shuffled datasets (shuffling was performed in a way to preserve the in- and out-degrees for each KN and PN respectively, as in Ref. [^43^] (Fig. 4G). We also found that the same glomeruli/ORs in the two flies reside near each other when placed in the PCA space of the first two dimensions (Fig. 4H). Our further analysis indicates that the first dimension in the embedding space is correlated with the sensitivity of olfactory receptors for food, while the second dimension has no clear functional significance (Supplementary Figure S1). Both the first and the second dimensions are not obviously related to the degree of connectivity of the KCs and instead are produced by glomeruli projecting to specific subsets of KCs (Supplementary Figure S2).

The first two *flat* dimensions of data explain only about 9% of connectivity data. As suggested by earlier work on the embedding of olfactory spaces^8,9^, the OR affinities may be better approximated by a curved low-dimensional manifold. To account for this possibility, we used Isomap algorithm^44^ (see Methods). The first two dimensions of the curved Isomap space account for about 39% (FlyEM) or 35% (FAFB) of the variance in the connectivity data (Fig. 4I, as quantified by the variance explained *within* the Isomap manifold). The datasets contain more variance along the first two dimensions (Fig. 6I) than shuffled data with preserved in- and out- degrees^43^. In addition, the individual glomerular connectivities, when embedded into a curved 2D manifold are situated in near locations in the two flies (Fig. 6H), suggesting that the structure of the affinity space is preserved between different individual animals within the same species. Overall, our findings suggest that the glomerulus-KC connectivity contains a low-dimensional structure as hypothesized in our primacy theory. Such connectivity structure can be identified in two different animals, suggesting that it is specified genetically. The structure is eliminated by the random shuffling of data which argues against the hypothesis of fully random OR-KC connectivity.

Is this conserved structure related to OR affinities for odorants? Above, we suggested that individual KCs should integrate inputs from OR displaying a high affinity for certain odorants (Fig. 3). This hypothesis, prediction (ii), can be tested directly by combining connectivity data (FlyEM, Fig. 5A) with the composite DoOR dataset^45^ (Fig. 5B). DoOR contains the most substantial description of Drosophila OR-ligand affinities available. Although not all OR-ligand pairs are present in the data (Fig. 5B), we can compute approximations of primacy sets for 156 odorants. To relate the odor response data to the connectivity data, we first performed an analysis of OR-OR similarity matrices in terms of their connections to KCs and in terms of their participation in the primacy sets. If KCs integrate inputs from high-affinity (primacy) ORs, the matrix of OR-OR similarities in connectivity (Fig. 5C) is expected to be correlated with the OR-OR primacy similarities (Fig. 5D). First, from the OR affinity matrix (5B left panel), we computed the primacy matrix for primacy number p=5 (Fig. 5B right panel), where each element is equal to one if a given OR belongs to the set of p=5 strongest responders to a given odorant. Then, using the primacy matrix we calculated OR-OR correlation matrix and compared it to OR-OR connectivity correlation matrix for the subset of glomeruli with a single projecting OR. We found that two matrices are significantly correlated (R=0.19, p<10^-4^), which suggests a correlation between high-affinity sets of receptors and OR-KC connectivity. The low value of the Pearson correlation is expected since we do not have access to the entire set of odorants ethologically relevant to flies and not all OR-ligand pairs are present in the data (Fig. 5B). The results of this analysis carried out for other values of the primacy number p=1-8 are shown in Supplementary Figure S5.

**Figure 5.**
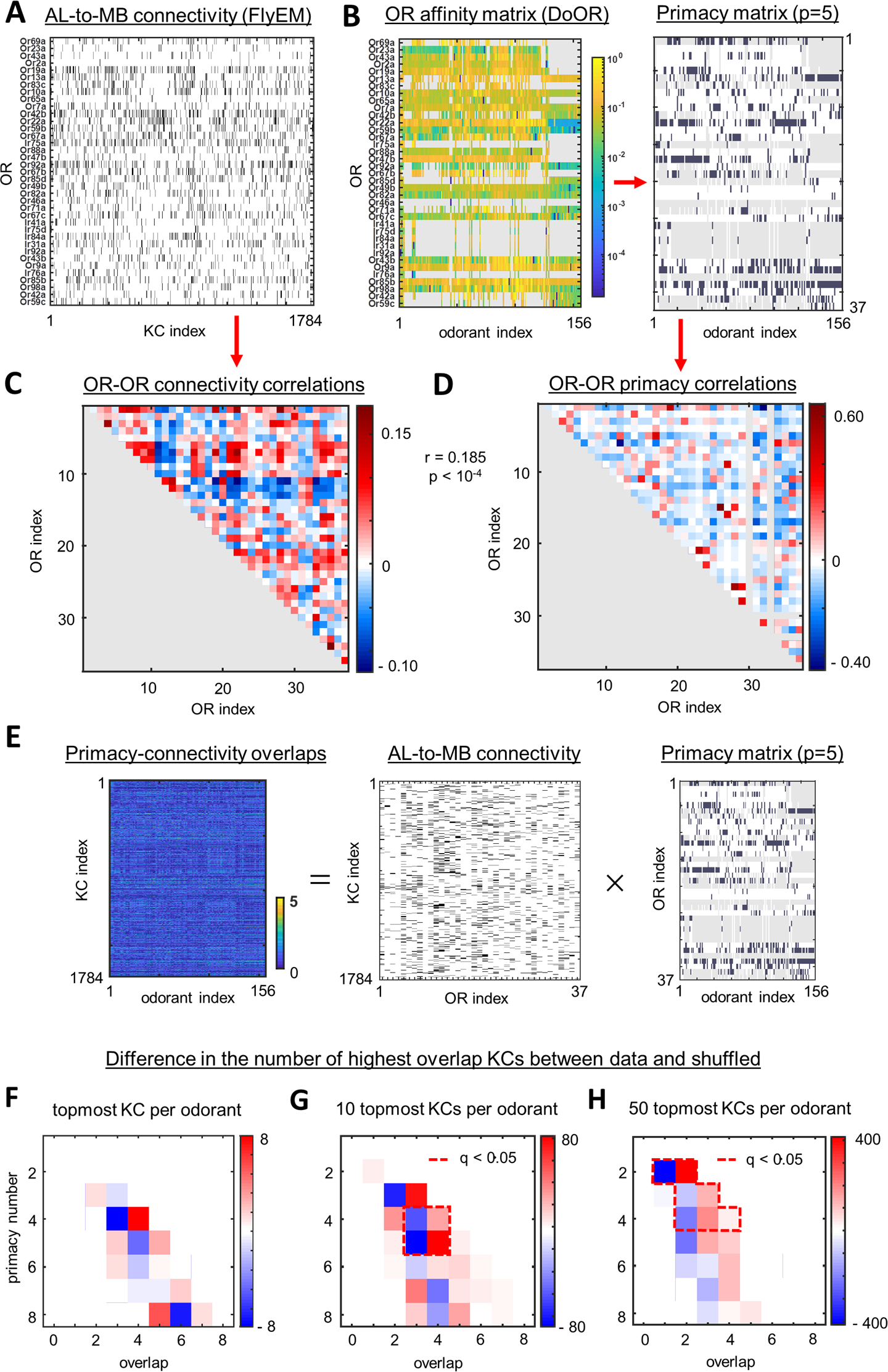
Comparing connectivity and affinity data. **A**. OR-KC connectivity matrix (FlyEM dataset). **B**. *Left:* OR-odorant affinities (from DoORv2 dataset). Gray represents missing values of affinities. *Right:* OR-odorant primacy matrix for p=5. **C**. OR-OR similarities (Pearson correlations) computed from connectivity data for 37 ORs represented in affinity data. **D**. OR-OR similarities (Pearson) computed from affinity data. The off-diagonal elements of matrices in **C** and **D** are correlated (R = 0.185, p < 10^-4^). **E**. Schematic showing the overlap matrix computed by matrix multiplication of connectivity and primacy matrices (p = 5). **F**. The difference in the number of actual overlaps of a given degree and the number of overlaps for a randomly shuffled connectivity matrix^43^. The difference is computed for different primacy numbers. Only the KCs with the highest overlap per odor (gKC) are considered. No statistically significant difference is observed. **G**. As in (**F**), but for the 10 topmost KCs per odor. Statistically significant differences are enclosed by a dashed red line (FDR<0.05). **H**. As in (G), but for the 50 topmost KCs per odor.

Can we directly compare the OR-KC connectivity matrices to the primacy sets of the odorants present in DoOR dataset? To perform this analysis, for each KC, we computed overlaps, (matrix product along the shared dimension) between the connectivity data and the primacy sets of ORs for each of the 156 odors present in the DoOR data. We can then find, for every odorant, a ‘grandmother’ KC (gKC), i.e. the KC which has the highest overlap between its connectivity with the primacy OR set for this odorant. Using this procedure, we can find 156 gKCs for each of the odorants present in the DoOR dataset and 156 corresponding overlaps. According to primacy theory, KCs should integrate inputs from primacy sets, so the overlaps between the gKC connections and the primacy sets are expected to be higher than for random connectivity. We find, however, *no* enrichment in the overlaps between connections and primacy sets for 156 gKCs identified in data compared to a randomly shuffled connectivity^43^ (Fig. 5F, FDR adjusted p-value (q-value) > 0.05). This finding suggests that the processing of the primacy information in the AL-MB network may not be based on the ‘grandmother’ KC mechanism (Fig. 3A, B).

Individual odorants activate >50 KCs in the MB^46, 47^, suggesting that a population of KC may represent different subsets of the primacy set for the odorants (Fig. 3C). We suggested that these subsets can be viewed as individual faces of the primacy simplex [Equation (6), Fig. 3C]. This mechanism is more robust than the one based on gKCs, in which only one cell is activated in the MB representing the primacy simplex. To test the population-based mechanism, for each odorant, we identified not one but 50 KCs with the highest overlaps between connections and primacy sets of ORs (Fig.5E). We thus evaluated 156 X 50 values of overlaps between connectivity and primacy sets for the odorants in the DoOR dataset. We have found a substantial enrichment in the number of higher overlap scores for real FlyEM connectivity compared to randomly shuffled connection matrices^43^ (Fig. 5H, FDR q<0.05). The enrichment can be seen in the presence of the red (positive) band to the right of the blue band in Fig. 5H indicating a larger number of higher overlaps compared to the random case. The enrichment in the overlaps between OR-KC connectivity and primacy sets is observed for the size of responsive KC population as small as 10 (Fig. 5G) for the primacy numbers in the range between 2 and 5. These findings indicate that, for individual odorants, a population of KCs has connectivity that is correlated with the primacy sets of ORs for this odor, rather than individual gKC. This correlation is significantly higher than that observed for randomized connectivity with preserved in- and out-degrees. Collectively, our analyses support our hypotheses regarding the processing of primacy information by the fly olfactory system by confirming i) the presence of a low-dimensional structure in feedforward connectivity that is shared across individual members of a species, and ii) that individual KCs integrate inputs from ORs with high affinity to an odorant (primacy sets).

## DISCUSSION

How can the nervous system link the representation of the same odorant at low and high concentrations? According to the primacy model, the odor identity is encoded by the OR types with the highest affinity for a given odorant, i.e., the primacy set of ORs. In air-breathing animals, odor exposure is defined by a sniff cycle, and the primacy set is activated at the beginning of a sniff cycle. As such, the primacy set is expected to be invariant to the ambient odor concentration and can link odor identity percepts across concentrations.

Hopfield proposed a model that attributes an odor identity to the sequence of receptor neuron activation ^48^. In his model, an increase in concentration leads to a temporal shift in the entire OR activity pattern^30, 49–51^. However, it is not clear how such a model can process the additional signals from receptors that were not activated at low concentrations but are recruited at higher concentrations. Alternative models of concentration invariant identity assignment based on the normalization of bulbar responses may only partially solve this problem ^52–55^. Indeed, normalization requires integration across all channels, or glomeruli ^56^ including those that are activated later in the sniff cycle^57^. Such mechanisms seem to preclude odor-guided decisions based on early olfactory inputs^25, 58^.

Several lines of experimental evidence support the primacy coding mechanism. To discriminate two odorants presented at random concentrations, animals use information accumulated during the short temporal window (∼100 ms) at the beginning of the sniff cycle^25^ and make these decisions before the entire ensemble of olfactory glomeruli is activated^57^. Similarly, the concentration-invariant cortical representations are found to be formed early in the sniff cycle^27^. A recent study, in which the timing of receptor activation was controlled optogenetically, demonstrated higher relevance of early activated glomeruli for sensory object identification^28^. Earlier studies in insects showed that the neural activity trajectories, in response to odor stimuli, diverge quickly for different odorants, but, initially, go together for the same odorant at different concentrations^59^. These results are consistent with the primacy coding mechanism.

The primacy model defines the primacy set as either the set of the most sensitive receptor types (affinity primacy) or as a set of the earliest activated receptor types/glomeruli (temporal primacy). In this study, we assumed that affinity and latency are highly correlated. While this is a reasonable assumption, OR activation latency may be affected by factors other than affinity, such as the solubility of an odorant in the mucosal layer or receptor distribution in the epithelium. Although the conclusions of the primacy theory are valid for both affinity and latency-based coding mechanisms, different forms of primacy may exist in different species. For example, in animals with a slow sniffing cycle, such as fish, affinity-based primacy may be a viable mechanism.

In addition to the *primacy coding hypothesis*, we proposed the hypothesis that the olfactory system samples a low-dimensional subspace in the space of odorant properties. We concluded that if olfactory space is low-dimensional and odorant identities are encoded according to a primacy code, evolutionary dynamics will drive ORs to reside along a thin high-affinity boundary that we call the primacy hull. In this model, those ORs that do not have a high affinity for any of the features of interest to the olfactory system will ultimately be pseudogenized. Thus, both the primacy and low-dimensionality assumptions are necessary for the primacy hull to exist. If, for example, the olfactory space is instead high-dimensional, the primacy mechanism may still be valid^25^. In this case, every OR can be a member of the primacy set, and, thus, evolution will favor the retention of all of the ORs in the genome. The hypothesis of the low-dimensional stimulus space is also compatible with some alternatives to primacy coding mechanisms. Primacy and low dimensionality are therefore two independent assumptions of our model.

Besides these two main assumptions, we have made many other simplifications that are frequently made in olfactory literature. For example, we assumed that receptor activation by odorant mixtures is given by a simple linear-nonlinear (LN) relationship [Equation (2)]. Although this approximation is conventional^35, 60, 61^, in light of significant recent work exploring the effects of non-linear interactions between mixture components^29, 30, 62–64^, we present an analysis of a more complex mixture model in Supplemental Materials S1. In the low concentration regime, in which the primacy sets are determined, we recover OR activations similar to equation 2, which justifies our use of LN approximation to describe primacy. The formation of the primacy hull is based on the assumption that most ORs that do not carry an olfactory function are eliminated in the course of evolution (pseudogenized). This assumption seems to be supported by the almost complete loss of ORs by aquatic mammals, such as dolphins and toothed whales^65^. If a substantial fraction of ORs were involved in alternative to olfactory functions, such a loss would not be possible.

Early identification of a predator or food source is a factor of evolutionary importance; by relying only on the highest affinity or shortest latency OR channels, a primacy code optimizes for the speed of percept formation. On the timescales of a single sniff, the primacy coding provides a quick and robust mechanism for identifying an odorant irrespective of the concentration at which it is encountered. An air-breathing animal may also increase the accuracy of its odor identification by sampling over several sniff cycles and integrating information to support a slower but potentially more accurate olfactory decision. This speed-accuracy tradeoff in olfaction has been previously explored^66^ and does not contradict the primacy mechanism, which operates on the time scale of a single sniff.

A primacy code involves the selective integration of early (primary) OR responses and can be neurally implemented via recurrent inhibition both in the OB^26^ and in the cortex^25, 27^. In the OB, the early activated glomeruli send signals to the cortex via mitral/tufted (MT) cells. Early MT cells activate an inhibitory bulbar network and may scramble or suppress the information from later activated MT cells^26^. Further in the cortex, early responses drive activity in a population of excitatory cells, which activate inhibitory interneurons via recurrent connections^27^. This results in ‘global’ inhibition in PC, which suppresses contributions from later responding (non-primary) ORs. This mechanism may implement a p-winner takes all (pWTA) circuit^25^. The primacy number could therefore reflect the average number of OR inputs necessary to drive global inhibition and produce a stable representation in PC or MB. The primacy number may not be fixed across different odorants. The number of OSNs and MT cells per OR type can vary widely^67, 68^ leading to differences in the excitatory drive provided by different OR channels. Assuming a fixed threshold for activating the global inhibitory shutdown of later responses, this would imply that the p-number for an odorant depends on the OR channels that it activates early in the sniff cycle. For example, if an odorant primarily activates ORs that are overrepresented at the OSN, glomerular, and MT levels at the start of the sniff cycle, one would expect the primacy number to be relatively low (lower than the average). The effective primacy number may also change if the cortical network structure results in different effective thresholds for different odorants or via adaptation of the number of OSNs per OR channel at the epithelium to sensory scene statistics^67^. In such cases, a stable odorant representation in PC/MB may still be achieved by pattern completion networks^69, 70^, which can compensate for degraded or incomplete inputs.

Our model makes two specific predictions regarding the structure of connectivity in the early olfactory system (AL to MB in insects or OB to PC cells in vertebrates). We proposed that primacy information may be processed if neurons in the target region (MB or PC) integrate inputs from neurons belonging to primacy sets. In this model, target neurons respond to activations of subsets of the primacy set (Fig. 3C), which makes the representations of odor identity in the target region robust to noise. Indeed, if a PC neuron corresponding to a particular subset of the primacy OR set is not activated, due to fluctuations in the inputs, this PC neuron may still be pushed over the activation threshold by the associative excitatory circuit in the PC. This prediction yields at least two corollaries. First, the low-dimensional structure of the odor space tessellated by the primacy sets should be present in the connectivity structure. As such, it can be revealed by a conventional dimensionality reduction method, such as PCA or Isomap. Second, the feedforward connectivity should be correlated with OR responses: PC/MB neurons tend to receive inputs from subsets of the primacy sets for specific odorants. We have tested these predictions using two recently obtained datasets on connectivity in the *D. Melanogaster* olfactory system^41, 42^ and the data on OR-odor affinities^45^. First, we found that connectivity data in two individual flies contains a similar low-dimensional structure. Second, we find that MB cells are more likely to receive connections from high-affinity (primacy) sets of receptors for individual odorants. These observations are consistent with the primacy coding hypothesis.

Recently, Zheng et al.^42^ characterized the structured component in the PN-to-MB connectivity matrix as deriving from a set of food-related glomeruli converging on KCs more frequently than would be expected under carefully constructed null models. This result is compatible with the proposed primacy hull-based connectivity model discussed here. First, this over-convergence is reflected in the loadings of glomeruli on the first principal component, where the identified food-related glomeruli have strong positive loadings, but are unrelated to loadings on the second principal component (Supplementary Figure 1). As both the first and second components are statistically significantly different from expectations under null models, the over-convergence of food-related glomeruli does not fully account for the observed structure in the connectivity matrix; the primacy hull may more fully account for this structure. Second, food-related glomeruli may have a wider role in the combinatorial coding of other not necessarily food-related odors; we may interpret these over-converging glomeruli as the important vertices/subsimplexes of the primacy hull, encoded in connectivity.

Overall, we have explored the implications of the primacy model^25^ which yields concentration-invariant odor identity representations based on the ORs most sensitive to a given odorant. We argue that evolutionary pressure to represent odorants according to a primacy code leads to the arrangement of OR types along a high-affinity surface called a primacy hull. Our presented analysis of fly olfactory datasets supports our predictions related to the implications of the primacy coding for olfactory circuit organization.

## METHODS

Our methods are described in Supplementary Methods online.

## ACKNOWLEDGEMENTS.

We thank Davy D. Bock for making fly connectivity data available prior to publication. This work was supported by the NINDS of the NIH under award number U19NS112953.

## Supplementary Information

### S1. Generating primacy hull in *D* dimensions P*_D_*, *_p_*

A large number of points, 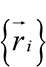 representing ORs, are distributed in the positive orthant in *D* dimensional space. F_→_ rom these, the points are selected via the following algorithm: i) generate a random unit vector 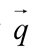 in the positive orthant, ii) compute 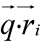 for all points indexed by *i* iii) select the *p* points with highest scalar product (the simplex) and add them to a list, *L*. Repeat steps (i)-(iii) until *L* does not change. The resulting *N* points (OR types) are the vertices of the primacy hull P*D*, *p*.

### S2. Generating simulated affinity data *K* containing

P *_N_* _,*D*, *p*_ : Given a primacy hull P *_N_* _,*D*, *p*_, vectors representing ORs can be collected into a matrix *R^N^*^×*D*^. Similarly, a panel of odorants represented by random unit vectors (always in the positive orthant) 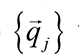 where *j* = 1,…, *N* can be collected as rows of matrix *P^D^*^×^*^Nodorants^*. Affinity data of dimensions *^N^odorants* ^×^*^N^* is simulated as *K* = *RP* (equation 4).

### S4. Linear model for OR responses to a mixture of odorants

Consider the mass-action law for an OR binding odorant number *o* :

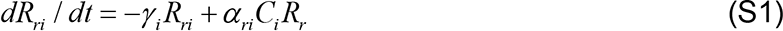

Here α *_ri_* and γ *_i_* are binding and unbinding rates for receptor *r* and ligand *i*, and the number of unbound (available) receptors of type *r* is given by

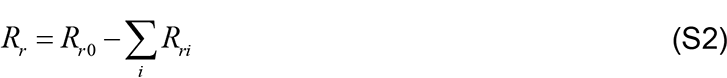

Here *R_r_* _0_ is the total number of OR molecules of type *r* exposed to odorant binding. Because in the equilibrium *dR_ri_* / *dt* and *R_ri_* = α*_ri_c_i_ R_r_* / γ *_i_* = *K_ri_c_i_ R_r_*, the total number of unbound receptors can be found from equation

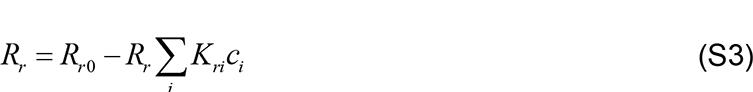

We thus obtain 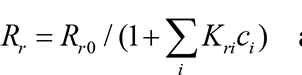 and the number of activated receptors

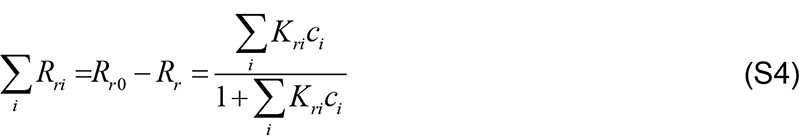

By assuming that the activity of the cell reflects the OR activation we obtain

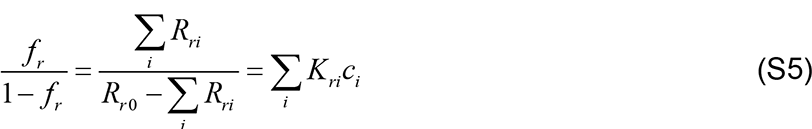

i.e. equations (1)-(2) of the main text.

### S5. Non-linear OR responses to mixtures and implications for the geometry of the primacy hull

In the main text we assumed that_→_an OR is activated if the following holds for its responses to a vector of concentrations *c _j_* = *cq_j_* 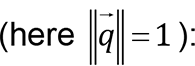:

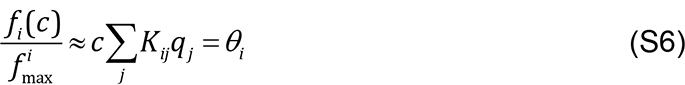

Here θ*_i_* = 1 is the threshold, for OR number *i*, at which this activation becomes noticeable to the system, and *c* is the overall concentration of the mixture. When applied to every OR, Equation (S6) allows us to determine both the set of primary receptors and the concentration of the mixture at which this set is fully activated. Here, we will identify the limits of applicability of this equation if the OR activation is determined by a more complex function that models mixture interactions.

A number or studies have shown that OR responses to mixtures of odorants are not always well captured by the linear-non-linear model used in the presentation of the primacy hull in the main text [1-5]. More complex models of OR-ligand dose response properties account for non-linearities in OR responses induced by mixture interactions, such as antagonism, synergy and masking [2, 4]. These models typically contain an additional parameter - η*_ij_* - which describes the *efficacy* with which a ligand *j* activates OR *i*. The general form for the activation of OR *i* by a mixture containing *M* components (obtained by integration of the rate equations from a two-step reaction model), is given by:

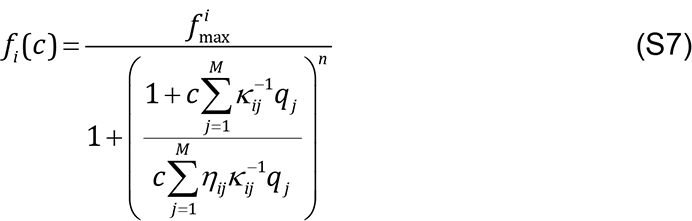

where the concentration of component *j* is given by *c _j_* = *cq_j_*, *j* runs from 1 to *M*, the number of monomolecular ligands in the environment and *q_j_* is the partial concentration of *j* in the mixture, *n* is the Hill coefficient and κ*_ij_* is the affinity of ligand *j* for OR *i*. This function can be approximated (for low concentrations of odorant) as:

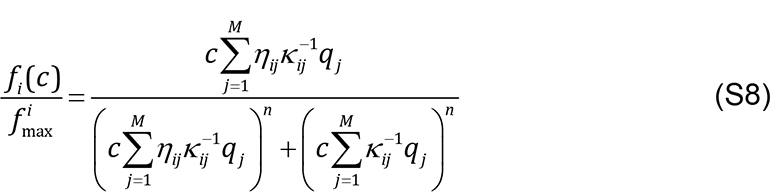

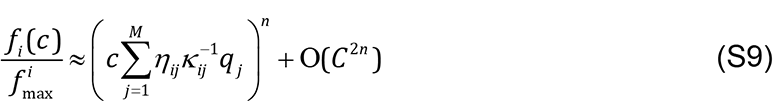

Those glomerular channels activated at the lowest mixture concentration to invoke a coherent percept constitute the primacy set of ORs. In order to elicit a downstream response in projection neurons, the firing rate of an OSN must exceed a threshold θ *_i_*. Dropping the second order terms in concentration, the relationship between affinity, efficacy and concentration at threshold is given by:

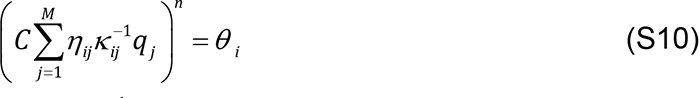

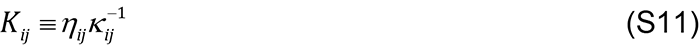

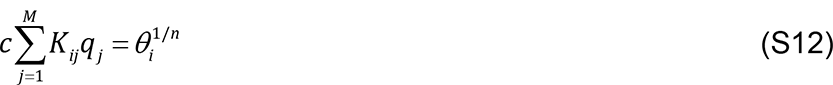

(S7) is of the same form as equation (S6), which was used in the main text to determine primacy sets. This suggests that our simple linear-nonlinear model can be used to determine primacy sets, even though OR activations are non-linear and include mixture interactions. This observation is based on the fact that primacy sets are determined at low odorant concentrations, at which the structure of OR response non-linearities is relatively simple and can be described by a single affinity matrix (S11).

The experimental literature documents cases in which Hill coefficients i) do not significantly vary across ORs or across OR-ligand pairs[6], ii) vary across ORs[7], iii) vary across odors for the same OR[7, 8]. In this latter case, where the Hill coefficient may be specific to each individual OR-ligand pair (ie. *n_ij_*), one theoretical study[9] suggests that the dose response curve of an OR-ligand mixture may be relatively invariant to manipulation of individual component Hill coefficients when compared to the other model parameters, at least in binary mixtures. We therefore assume equation S1 to be a good approximation to Equation S7 and sufficient to define the primacy set for an odorant mixture.

This approximation tells us that the higher the value of *K_ij_*, the lower the mixture concentration at which the response of OR *j* meets the threshold for evoking a downstream response and relaying partial information about the odorant. If the relevant space in which to consider the evolution of the OR repertoire was *K* -space in the main text, we propose that within this framing of receptor responses, ORs instead evolve in *A* -space. For fixed threshold θ *_i_*, the product of affinity and efficacy defines the coordinates of an OR in this space. Assuming a primacy code for the computation of odor identity and a given environmental odor statistics, individual ORs will evolve to balance their affinities and efficacies to a subset of the physico-chemical properties of interest to the olfactory system. This balance will involve maximizing the product of these variables subject to constraints imposed by the combinatorial nature of the primacy code, which requires ORs to respond with similar frequencies to the odor environment for an optimal code capacity[10]. We suggest that evolutionary optimization via e.g. a birth death-model[11] will drive the OR repertoire to be distributed as a primacy hull in *A* -space over an evolutionary timescale.

### S6. Formatting connectivity and affinity data: *FlyEM and FAFB*

Each uPN-KC connectivity matrix *C*_*PN*_ was first reduced to a glomerulus-KC connectivity matrix *C*_*GL*_ by considering each glomerulus label in turn, summing all uPN connectivity vectors (rows in *C*_PN_) associated to that glomerulus and adding the resulting vector as a row in the binary matrix *C*_GL_. *List of olfactory glomeruli:* D, DA1, DA2, DA3, DA4l, DA4m, DC1, DC2, DC3, DC4, DL1, DL2d, DL2v, DL3, DL4, DL5, DM1, DM2, DM3, DM4, DM5, DM6, DP1l, DP1m, V, VA1d, VA1v, VA2, VA3, VA4, VA5, VA6, VA7l, VA7m, VC1, VC2, VC3, VC4, VC5, VL1, VL2a, VL2p, VM1, VM2, VM3, VM4, VM5d, VM5v, VM6, VM7d, VM7v. A reduced version of this matrix was used for the analyses, shown in Fig.5, with OR labels corresponding to those in the DoOR affinity dataset. *List of subset of glomeruli:* D, DA3, DA4l, DA4m, DC1, DC2, DC3, DL1, DL3, DL5, DM1, DM2, DM4, DM6, DP1l, VA1d, VA1v, VA2, VA3, VA4, VA5, VA6, VA7l, VC2, VC4, VC5, VL1, VL2a, VL2p, VM1, VM2, VM3, VM4, VM5d, VM5v, VM7d, VM7v, with the corresponding ORs determined as shown in Figure S4 and listed below.

#### DoOR OR-odorant affinity

We extracted a dense subset of affinity scores from the DoOR dataset[12]. This incomplete matrix contains affinities, which take values in the interval [0, 1], for 3488 OR-odorant pairs (60% complete). The resulting matrix is of dimension 37 (ORs) x 156 (odorants). We selected ORs i) that have a clear cognate glomerulus, i.e., all inputs to the glomerulus come from OSNs exclusively expressing that OR, and ii) for which there is sufficient data to distinguish relatively high affinities and define approximate primacy sets, i.e. ORs for which there are at least 18 (2 max(*P*) + 2) odorant affinities recorded. *List of glomeruli in dense subset of DoOR:* Or69a, Or23a, Or43a, Or2a, Or19a, Or13a, Or83c, Or10a, Or65a, Or7a, Or42b, Or22a, Or59b, Or67a, Ir75a, Or88a, Or47b, Or92a, Or67b, Or85d, Or49b, Or82a, Or46a, Or71a, Or67c, Ir41a, Ir75d, Ir84a, Ir31a, Ir92a, Or43b, Or9a, Ir76a, Or85b, Or98a, Or42a, Or59c.

### S7. Comparing FlyEM and FAFB datasets along the PN dimension via Monte Carlo alignment

To compare the binarized PN-KC connectivity matrices from the FlyEM and FAFB datasets (*W*^*FlyEM*^_*PN,KC*_ and *W*^*FAFB*^_*PN,KC*_ respectively), we aligned them along the unlabeled KC dimension as follows. First, we computed a distance matrix between the KCs belonging to different datasets *D_FlyEM,FÆFB_*. For each pair of the KCs, the distance between them was defined as one half of the squared count of mismatching inputs from the PNs minus the squared count of matching inputs from the PNs:

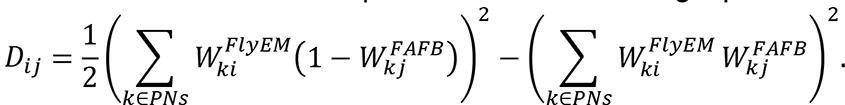

This way, the matches were prioritized compared to the mismatches, reflecting the bias in the data annotation process where only the most certain projections were included in connectivity matrices. We then coarsely aligned the KCs of the two datasets. To do that, we initialized an empty matrix *W*^∗^_*PN,KC*_ of a size of the larger (FlyEM) dataset. Then, on every iteration, we selected a random KC *i* with no repetitions from the larger (FlyEM) dataset and used the distance matrix *D* to determine the most similar KC *j* from the smaller (FAFB) dataset: 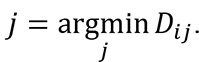. The projection pattern *W*^*FAFB*^_*PN,j*_ of the KC *j* was then removed from the smaller (FAFB) dataset and pasted to the matrix *W*^∗^_*PN,KC*_ at the position *i* matching that of the selected KC from the larger (FlyEM) dataset: *W*^∗^_*PN,KC*_ = *W*^FAFB^. The PN,i PN,j process repeated until no KCs remained in the smaller (FAFB) dataset. Finally, to improve the match between the larger dataset *W*^FlyEM^ and the coarsely aligned smaller dataset *W*^∗^_*PN,KC*_, we permuted the latter along the KC dimension using a greedy Monte Carlo algorithm. Specifically, on every iteration, we permuted two random KCs *i*, *j* in the coarsely aligned smaller dataset *W*^∗^_*PN,KC*_ and computed the resulting change in distances: Δ*E* = *D*_ii_ + *D*_jj_ — *D*_ij_ — *D*_ji_. If this change Δ*E* was less than zero, we accepted the permutation; otherwise, we rejected it. We performed a total of 10^7^iterations of the algorithm. To evaluate the alignment, we computed the fraction *f* of synapses in the aligned smaller (FAFB) dataset matching the synapses of the larger (FlyEM) dataset: *W*^∗^_*PN,KC*_ = *W*^FlyEM^. We evaluated the algorithm 10 times for the (same) data matrices and also 10 times for the (different) null models where both marginals were preserved. We then compared the alignment qualities *f* for the data and null models using the paired-sample t-test with a threshold p-value of 10^−3^.

### S8. Comparing FlyEM and FAFB datasets along the OR dimension by analysis of correlations

To compare the binarized PN-KC connectivity matrices from the FlyEM and FAFB datasets regardless of the KC order, we performed the correlation analysis as follows. For each PN-KC connectivity matrix, we computed a matrix of Pearson correlations between the PN projection patterns: *W_PN,KC_* → *C_PN,PN_* where *C*_ij_ = *corr*(*W_i_*_∗_, *W_j_*_∗_). We then computed the Pearson correlations between these matrices for different datasets: ℭ = *corr*(*C^FlyEM^*, *C^FAFB^*). We performed this procedure 10 times for (different) bootstrapped versions of the two datasets (where the KCs in each dataset were selected randomly with repetitions) and also 10 times for the (different) null models where both marginals were preserved. We then compared the correlation coefficients ℭ for the bootstrapped data and null models using the paired-sample t-test with a threshold p-value of 10^−3^.

### S9. Comparing FlyEM and FAFB datasets along the OR dimension via dimensionality reduction

To analyze the dimensionality of the binarized PN-KC connectivity matrices from the FlyEM and FAFB datasets, we used the dimensionality reduction techniques as follows. First, we normalized the binary connectivity matrices by shifting the means of every PN projection pattern to zero and scaling their variances to one. To obtain linear embeddings of the data, we performed the principal component analysis (PCA) on normalized projection matrices and computed the per-dimension variance of the PCA representation. To obtain non-linear embeddings of the data, we performed Isomap on normalized projection matrices and computed the per-dimension variance in the Isomap space in the same way with the variance computed in the previous step. To determine the optimal number of nearest neighbors to be used in the Isomap algorithm, we performed Isomap with the numbers or nearest neighbors from the minimum of 2 to the maximum of 50 using both FlyEM and FAFB datasets. For each number of nearest neighbors *k*, we computed the L2 norm of the difference Δ between Isomap representations of FlyEM and FAFB considering only the two first Isomap dimensions. We determined the optimal number of Isomap dimensions (*k* = 6) by locating the elbow of the Δ(*k*) plot where the difference Δ began saturating as a function of the number of nearest neighbors *k*. We performed PCA and Isomap procedures for FlyEM and FAFB datasets, as well as for the corresponding (different) null models, 10 for each dataset. For the null models, we computed the mean and the variance of the pre-dimension variance explained.

### S10. Analysis of KC overlaps (**Figure 5**)

To compare the overlaps between OR-KC connectivity data and primacy sets contained in the OR-odorant affinity data, we first compute the overlap matrix (Fig. 5F) via the product *O*(*p*) = *AS*(*p*). Each entry *O*(*p*)*_ko_* represents the overlap between two binary vectors – one the OR inputs to KC number *k*, *A_k_* _,._, and the other, the primacy set corresponding to odorant *o*, *S* (*p*)_.,*o*_. For a given primacy number *p*, the matrix elements contained in *O*(*p*) can be summarized as a histogram. First, we identified the overlaps for the ‘grandmother’ KC for each smell: the strength of connectivity of the highest overlapping KC with the corresponding primacy set. This strength can be combined in the vector *h*(*p*,1)*_o_*, which contains the amount of overlap for odor *o* between its primacy set and the best, ‘grandmother’ KC, for each primacy number. The histogram of this vector is shown in Figure 5F (minus the null model). Next, we computed the strengths of overlap for *l* topmost KCs (*l* = 1) for the case of ‘grandmother’. This yielded the quantity *h*(*p*,1)*_o_*. This quantity is actually a tensor, which, for each odorant *o*, and a primacy number *p*, contains *l* topmost overlaps. The histograms of values of this object are shown in Figures 5G and H, with the null model subtracted.

We compare the empirical distribution of overlaps contained in *h*(*p*,1)*_o_* to the null model in which correlations are removed from the connectivity data by shuffling, we repeat the above analysis for each shuffled connectivity matrix, yielding an ensemble of matrices. We report the difference between empirical and null distributions via the difference matrix. Positive values (shown in red in Fig. 5F-H) indicate that there are more overlaps of a given degree between the empirical connectivity and primacy matrices, corresponding to sub-simplexes or faces of primary simplexes, relative to the null distribution. Negative values (shown in blue in Fig. 5F-H) indicate that the data contains fewer overlaps than would be expected under our null model. Statistical significance is assessed by computing an FDR corrected p-value for each tuple [(p, overlap)].

## Supplementary Figures

**Figure S1.**
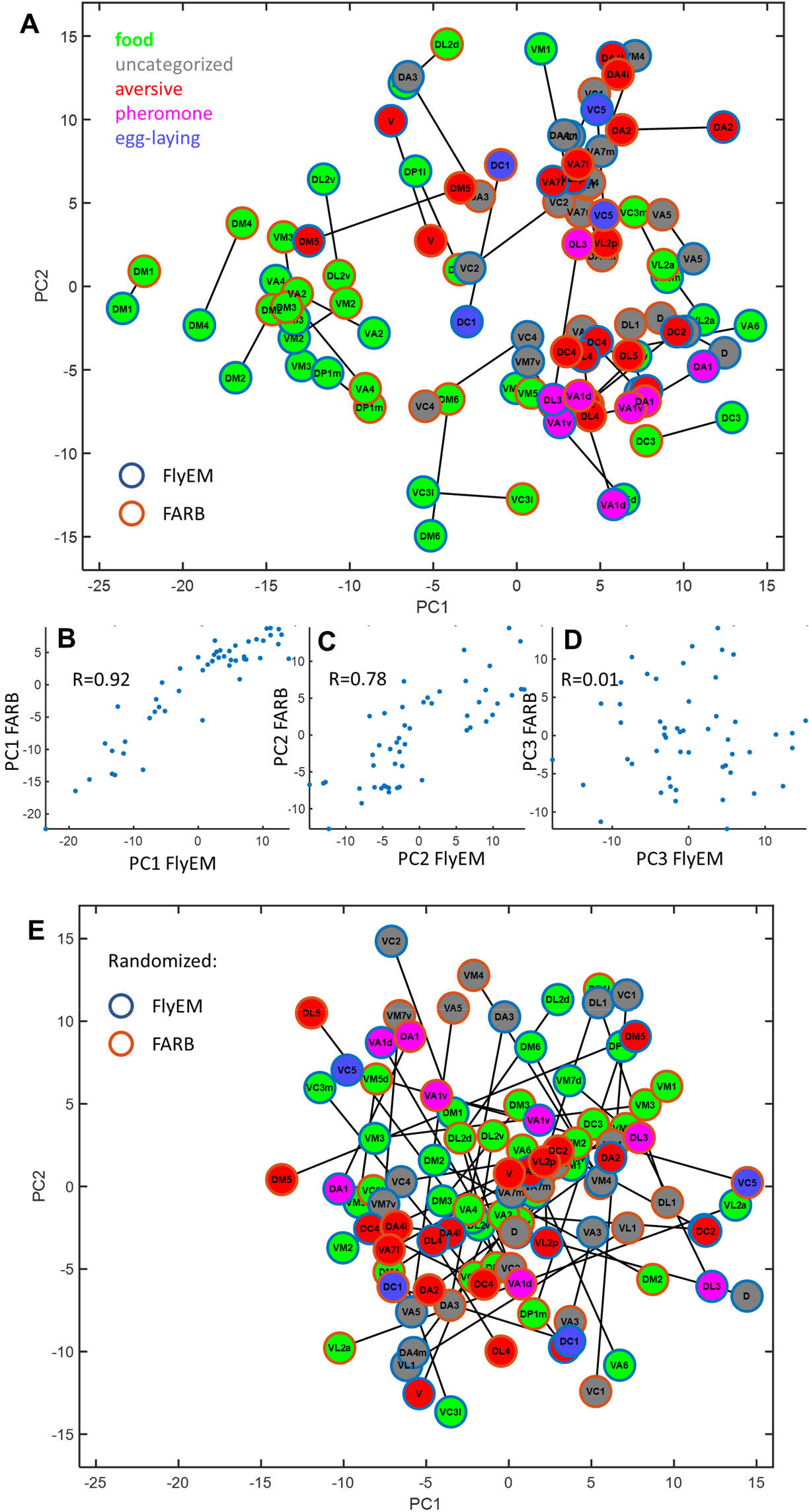
Functional significance of non-random features of connectivity in FlyEM and FAFB datasets. (A) Glomeruli placed in the connectivity’s 2D PCA space. The same glomeruli in two datasets are connected by lines. Glomeruli are colored according to their function as indicated. The first PC of connectivity appears to be related to food-sensitive glomeruli, while the second PC is unrelated to food. (B-D) Three first PCs for the two datasets plotted against each other. The first two PCs (B and C) are conserved between FlyEM and FAFB datasets (R=0.92 and 0.78), while the third PC appears to be random. This indicates that only first two PCs of connectivity are conserved across individual animals. (E) For randomly shuffled connectivity matrices, none of the principal components are conserved across individuals.

**Figure S2.**
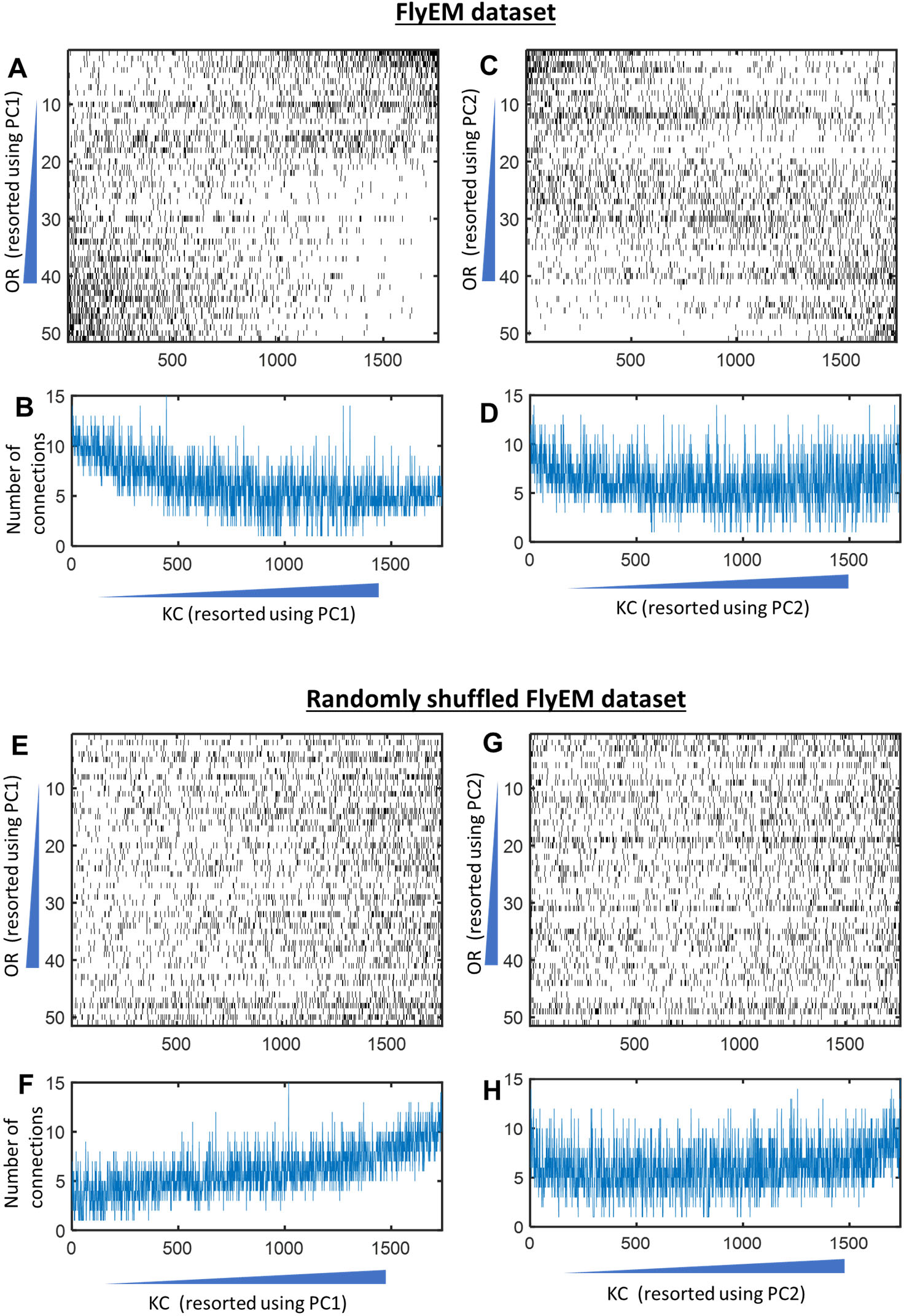
The first two PCs of connectivity cannot be explained by in-degree of the KCs. Instead, they are related to the connectivity structure. (A) Binarized connectivity matrix in FlyEM dataset with both ORs and KCs sorted according to their contribution to the first PC (PC1). ORs with similar PC1 appear to have stronger connectivity, suggesting that the structure of OR-KC connections determines the contribution of ORs to a PC. (B) the number of connections made by KC in the binarized matrix does not have a clear monotonic dependence on PC1. Thus, the first PC is not produced by differences in KC in-degree. (C, D) Same for PC2. A diagonal band along the diagonal in the sorted connectivity matrices in (A) and (C) indicates that ORs are connected to specific groups of KC, which determines both PC1s. Thus PC1 and PC2 emerges from a specific connectivity structure. (E-H) The same analysis in the in randomized matrices shows that the first PC is correlated with the KC in-degree (F).

**Figure S3.**
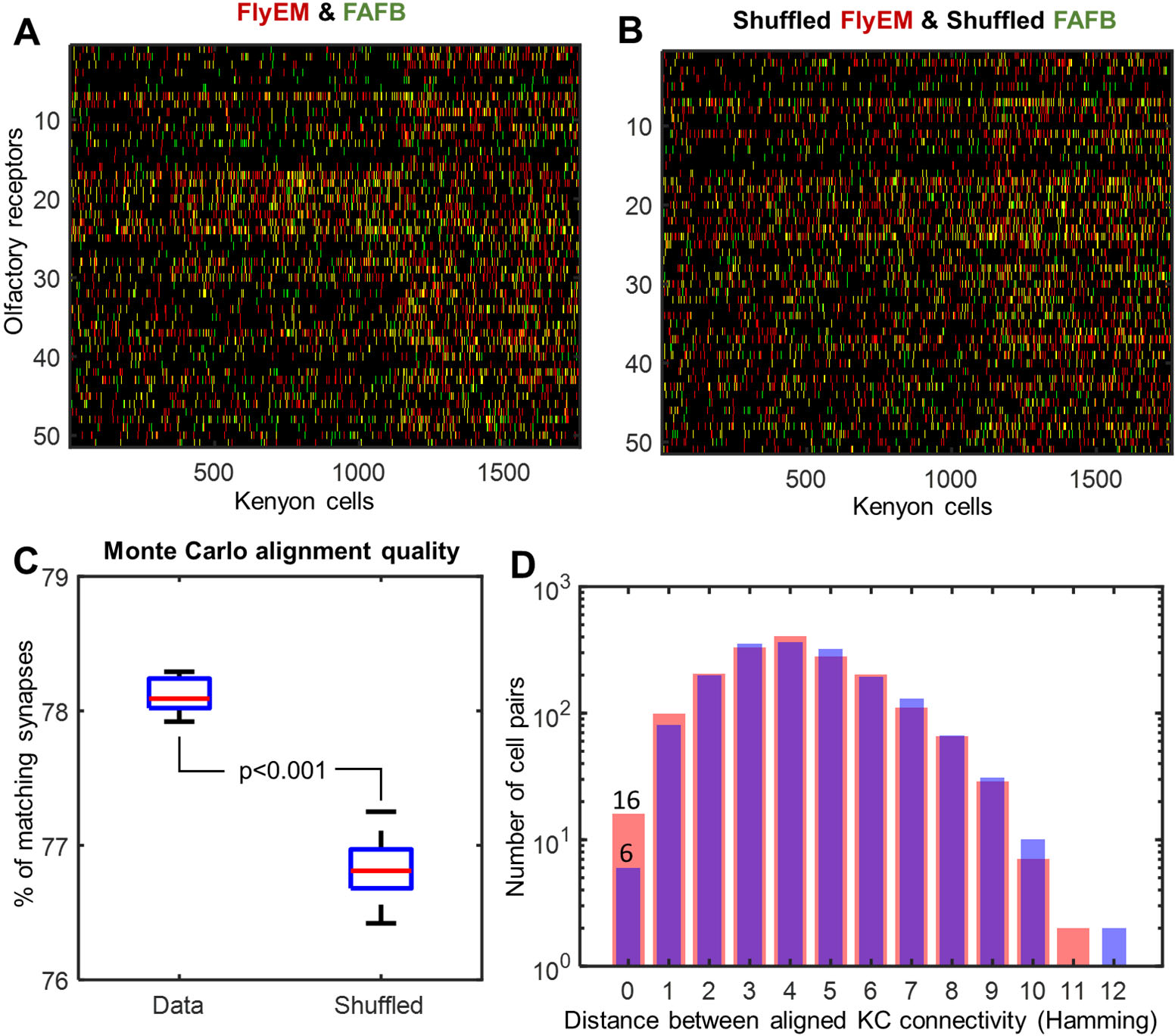
Results of brute force alignment of connectivity datasets. (A) Two connectivity datasets aligned using the simulated annealing algorithm. (B) Same for two randomly shuffled connectivity matrices. (C) Even randomly shuffled connectivity matrices share ∼77% of synapses, when aligned. Unshuffled connectivity matrices share ∼78% synapses. (D) Hamming distances between aligned KCs. Only 16 KCs are an exact match (H=0) versus 6 KCs in the randomly shuffled case.

**Figure S4.**
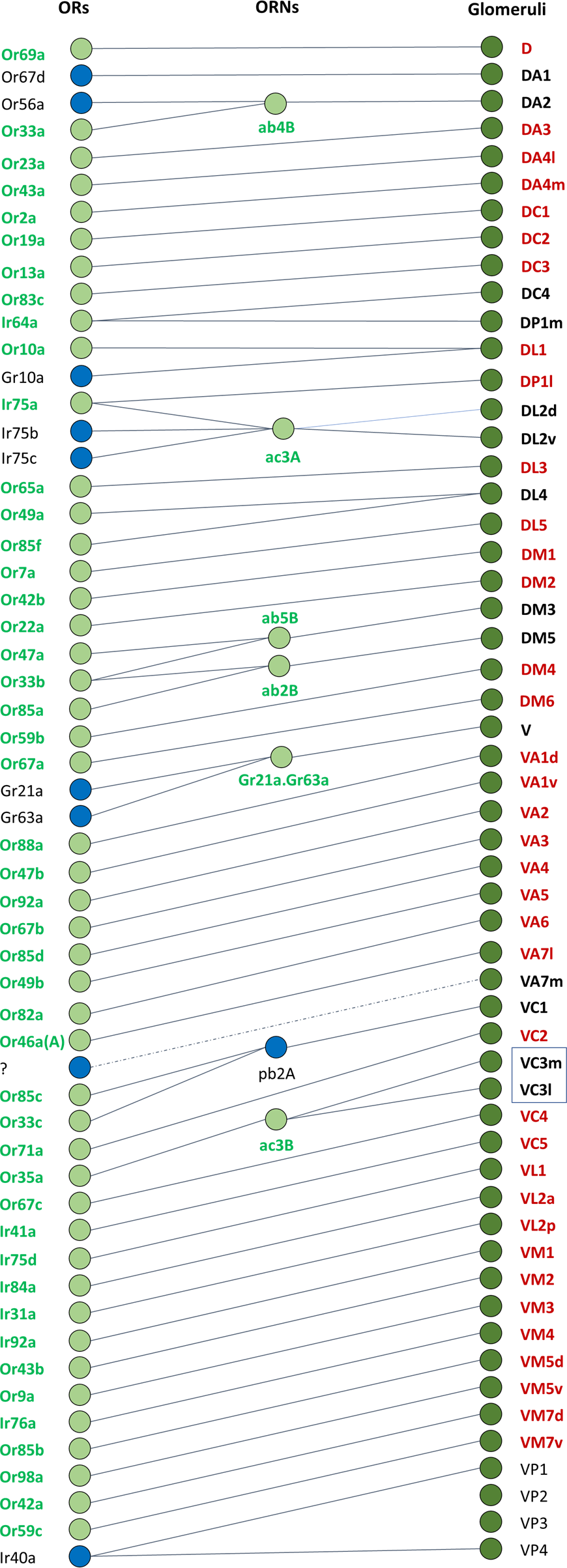
Glomeruli and their cognate odorant receptors. Each glomerulus in the *Drosophila* antennal lobe receives input from specific OR types. The majority receive input from only a single odorant responsive receptor type, however some glomeruli, e.g. DL1, receive converging input from >1 OR type. *Left:* OR types present in the DoOR dataset are represented by a filled green circle, while a blue circle represents those ORs missing from the DoOR. *Right:* Glomeruli that are included in Figures 4 and S1 are labelled in bold and those that are included into our analysis of DoOR data (Figure 5) are labelled in bold red. Based on data from [13].

**Figure S5.**
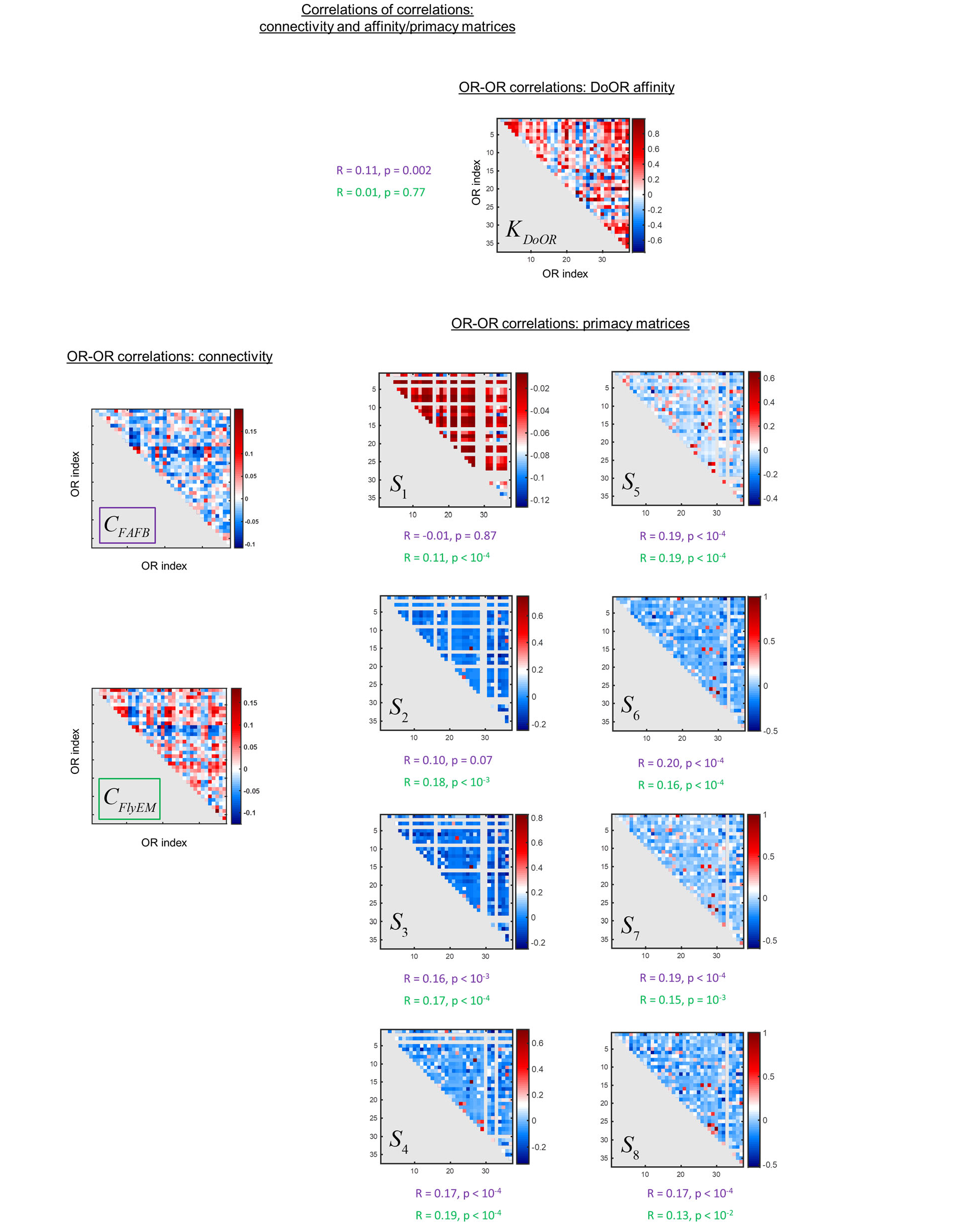
*Left:* OR-OR Pearson correlation matrices for FlyEM and FAFB connectivity data (37 glomeruli). *Right*: OR-OR Pearson correlation matrices for primacy sets in DoOR affinity data shown for a range of primacy numbers. Correlations between connectivity and primacy sets are reported for FlyEM (in purple) and for FAFB (in green). Statistically significant correlation coefficients are recorded for a range of primacy numbers.

## Notes

### Competing Interest Statement

The authors have declared no competing interest.

### Summary of Updates

Edited for clarity and shortened. Tests of the primacy model using fruit fly data are added.

